# Evolutionarily conserved ovarian fluid proteins are responsible for extending egg viability in salmonid fish

**DOI:** 10.1101/2023.09.05.556347

**Authors:** Aurélie Gueho, Daniel Żarski, Hélène Rime, Blandine Guével, Emmanuelle Com, Régis Lavigne, Thaovi Nguyen, Jérôme Montfort, Charles Pineau, Julien Bobe

## Abstract

In contrast to most fish species, salmonids exhibit the unique ability to hold their eggs for several days after ovulation without significant loss of viability. During this period, eggs are held in the body cavity in a biological fluid, the coelomic fluid (CF) that is responsible for preserving egg viability. To identify CF proteins responsible for preserving egg viability, a proteomic comparison was performed using 3 salmonid species and 3 non-salmonid species to identify salmonid-specific highly abundant proteins. In parallel, rainbow trout CF fractions were purified and used in a biological test to estimate their egg viability preservation potential. The most biologically active CF fractions were then subjected to mass spectrometry analysis. We identified 50 CF proteins that are overabundant in a salmonid-specific manner and present in analytical fractions exhibiting the highest egg viability preservation potential. Here we show that salmonid CF is a complex biological fluid and that several proteins are responsible for preserving egg viability. Among these key players are proteins related to immunity, calcium binding, lipid metabolism, proteolysis, extracellular matrix and sialic acid metabolic pathway.

## Introduction

In most teleost fish species, egg viability and quality (*i.e.*, the egg ability to be fertilized and subsequently develop into a normal embryo) decreases rapidly after ovulation ^1,2^. This is for instance the case in goldfish (*Carassius auratus*), in which a decrease in egg quality is detected as early as 3 hours post-ovulation (hpo) at 23°C and a complete loss of viability is observed 12 hpo ^3^. In common carp (*Cyprinus carpio*), 50 % of the eggs lose their viability within 6 hours after ovulation, and the complete loss of viability is reached 12 to 14 hpo ^4^. In sharp contrast with all of other teleost fish species, salmonids have the unique ability to hold their eggs for several days, or even weeks, following ovulation without any significant loss of their viability ^5–7^. In contrast to most teleost fish, salmonid fish exhibit “open” ovaries (termed secondary gymnovarians) that are devoid of ovarian lumen. At ovulation, eggs are therefore released directly in the body cavity where they bath in a semi-viscous fluid, called coelomic fluid (CF), or sometimes ovarian fluid (OF) in consistency with other teleost species that exhibit a closed ovary and because some OF components are of ovarian origin^8–10^.

Several studies have shown that salmonid egg viability and ability to be fertilized decreases rapidly after ovulation when eggs are held *in vitro* in artificial medium rather than in CF ^11^. This indicates that CF components plays a major role in maintaining egg viability. In addition to its unique ability to maintain egg viability both in vivo and in vitro, CF also exhibits several features associated with fertilization. In fresh water, eggs are water-activated within a minute. The micropyle (*i.e*., the canal through which the spermatozoa penetrates into the oocyte during fertilization) closes and the chorion hardens making fertilization impossible ^12–14^. As CF is expelled along with the eggs into fresh water during spawning, its presence around the eggs, even diluted, is believed to locally contribute to maintaining an optimal environment able to prevent or delay immediate egg activation to allow fertilization ^15^. CF would also play a chemoattractant role during fertilization to guide the spermatozoa to the micropyle when male and female gametes are simultaneously released into the water ^16^. In addition, salmonids coelomic fluid can also be used to maintain egg viability in non-salmonids species like goldfish ^17,18^ or zebrafish ^19^.

The physicochemical properties of salmonid CF have been extensively studied. CF pH is around 8.5 and its osmolarity is about 290 mmol.kg^-1^. The ionic composition of CF is similar to blood plasma. It contains sodium (Na^+^), potassium (K^+^), calcium (Ca^2+^), magnesium (Mg^2+^), chloride (Cl^-^) with differences in the amount of K^+^ as CF contains 3-4 times more K^+^ than blood plasma. CF also contains glucose, fructose, proteins and free amino acids, cholesterol and phospholipids ^20–22^. As indicated above, several lines of evidence strongly suggest that mimicking the ionic composition of the trout CF is not sufficient to fully preserve egg viability. This also indicates that the unique biological features of salmonid CF are conferred by organic compounds and most probably by proteins. While the general physicochemical composition of salmonids CF is well known, our knowledge of CF proteome is, in contrast, extremely limited. CF is known to have different enzymatic activities like phosphatase, collagenase, gelatinase and lactate deshydrogenase activities ^20^. Several CF proteases have been identified including a progastricsin ^10^ and a serine protease of the chymotrypsin family^23^. Some protease inhibitors, named Trout Ovulatory Proteins (TOP) were also identified in brook trout (*Salvelinus fontinalis*) and rainbow trout (*Oncorhynchus mykiss*) CF ^8,9^. The TOP proteins have the particularity to confer anti-bacterial activity to the CF. Using 2D proteomic analysis, potential markers of egg quality, such as vitellogenin fragments and lipoproteins that may originate from leaking eggs and be the consequence of oocyte postovulatory ageing in CF have been reported ^24^. The presence of components from broken eggs was found to decrease the pH of trout CF ^25^ indicating that the biochemical composition of salmonids CF can indirectly reflect the quality of the eggs in the case of post-ovulatory ageing. More recently, shotgun proteomic analysis of rainbow trout CF ^26^ and of chinook salmon CF ^27^ led to the identification of 54 and 174 proteins, respectively. Despite this, our current knowledge of salmonid CF proteome is still partial and proteins responsible for its biological properties remain unknown.

The aim of this study was to comprehensively and functionally characterize proteins of rainbow trout CF and subsequently identify the proteins responsible for the extension of egg viability in salmonid CF. For this, the proteomic composition of different salmonid CF and non-salmonid OF was compared to identify proteins specifically present in salmonids CF. We reasoned that salmonid-specific proteins could be responsible, at least in part, for preserving egg viability, a feature found only in salmonids. In parallel, rainbow trout CF was fractionated using HPLC coupled to gel filtration or ion exchange columns in order to subsequently identify protein fractions with strong egg viability preservation potential. Proteins present in the fractions allowing the highest egg viability preservation were identified using mass spectrometry. Identifying the CF components involved in preserving egg viability would have important applied outputs with various biotechnological applications for aquaculture. It could also help understanding egg quality preservation mechanisms, and subsequently improving egg storage condition in salmonid and non-salmonid fish species.

## Results

### Proteins are responsible for the biological activity of rainbow trout CF

Rainbow trout eggs were stored in different dilutions of rainbow trout CF (100%, 75%, 50%, and 25%) diluted in mineral medium (MM) mimicking the CF mineral composition^28^ or in MM only. I*n vitro* fertilization was carried after 3 days at 12 °C to assess the impact of storage on egg viability and ability to be fertilized. No significant differences were observed in the percentage of eggs reaching eyed-stage between eggs stored in the different CF dilutions and eggs stored in the non-diluted CF. The survival rate (*i.e*., embryos reaching eyed stage) was however dramatically reduced when eggs were stored in mineral medium alone (Figure 1A). These observations indicate that organic (*i.e.*, non-mineral) components of the CF are responsible for maintaining egg viability and ability to be fertilized for several days after ovulation. Our observations also suggest a strong biological activity of at least some of the non-mineral CF components as a 25% CF dilution was sufficient to maintain an egg viability similar to that observed using non-diluted CF.

**Figure 1.**
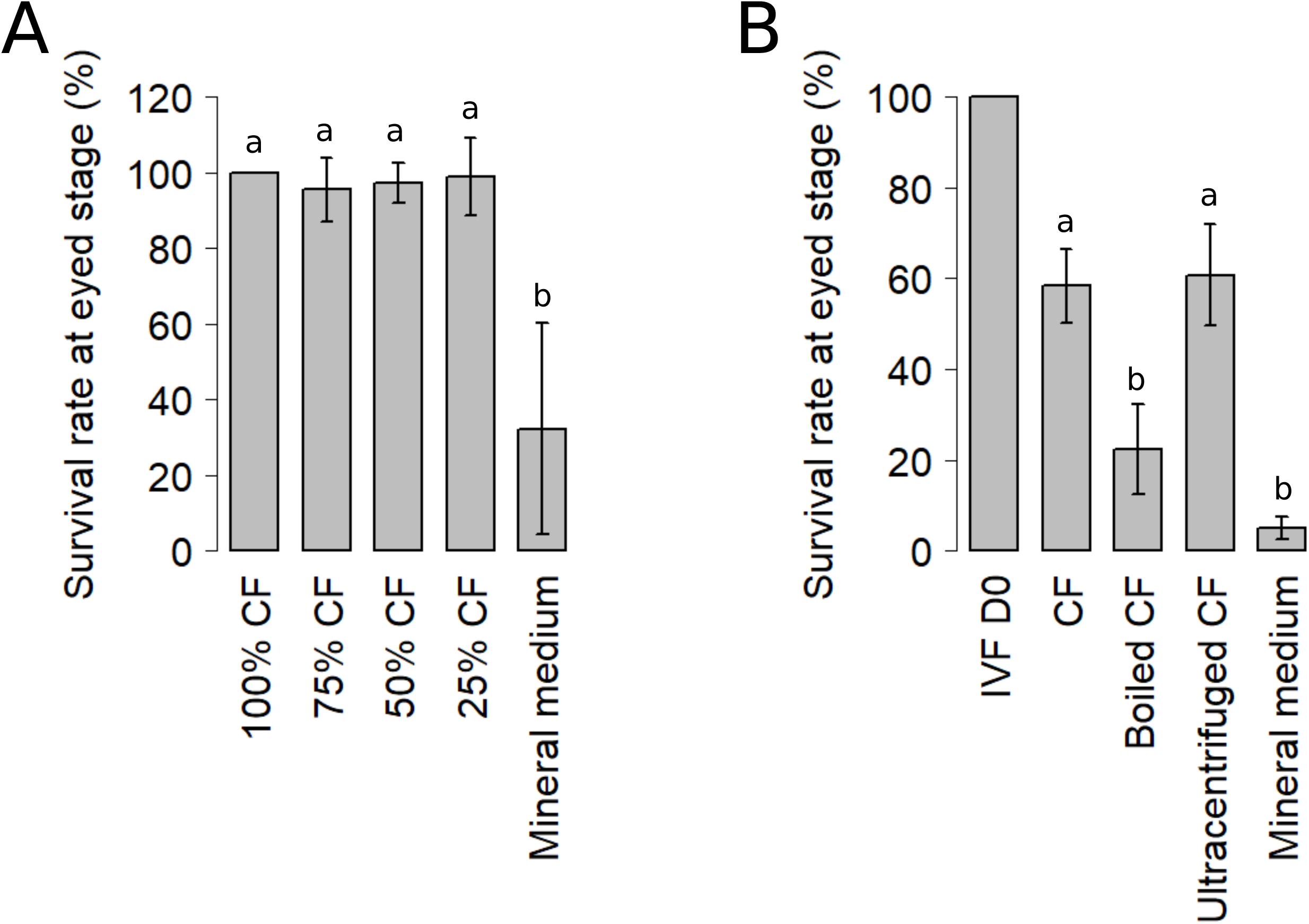
Effect on eying rates of 3 days of in vitro storage of unfertilized rainbow trout eggs in different dilutions of CF (A). Results have been normalized to the percentage of eggs reaching eyed stage in the non-diluted CF condition (100% CF). Percentages of eggs reaching eyed stage (mean±sd, n=5) superscripted with the same letter are not significantly different, Wilcoxon-Mann Whitney test (p-value<0.05). Effect on eying rates of a 3 day in vitro storage of unfertilized rainbow trout eggs in coelomic fluid (CF), boiled coelomic fluid (Boiled CF), ultracentrifuged coelomic fluid (Ultracentrifuged CF) or trout mineral medium (B). Results have been normalized to the percentage of eggs reaching eyed stage when the fertilization is done immediately upon eggs collection (IVF D0). Percentages of eggs reaching eyed stage (mean±sem, n=7) superscripted with the same letter are not significantly different, t-test (p-value<0.05).

To identify the nature of the organic compound conferring its biological activity to the trout CF, eggs were stored for 3 days at 12 °C in CF (*i.e*., non-diluted), boiled CF, ultracentrifuged CF, or MM before fertilization. CF was boiled to denature the proteins while ultracentrifugation was used to remove the lipids in order to test the respective contribution of proteins and lipids. No significant difference was observed in the percentage of eggs reaching eyed stage between eggs stored in CF and eggs stored in ultracentrifuged CF. However, when CF proteins had been denatured before holding eggs, this percentage was significantly decreased (Figure 1B). This suggests that proteins, but not lipids, are responsible for the biological activity associated with CF ability to maintain egg viability and ability to be fertilized.

### Proteomic characterization of rainbow trout CF

In order to understand how rainbow trout CF allows egg conservation, its proteomic composition was characterized by mass spectrometry analysis. Together, 573 proteins were identified (Table S1). A large majority of the proteins were newly identified in the trout CF compared to previous proteomic studies ^26,27^ allowing the characterization of a more complete trout CF proteome. Human orthologs could be found for 482 of those proteins, corresponding to 326 unique human genes. Those 326 unique genes were then used for biological processes GO-term annotation and the 500 most enriched GO-terms found were clustered into several processes (Figure 2A, Table S2). Fifteen main clusters connecting at least 4 different GO-terms sharing 70% of their genes were identified. For each of them, the corresponding high-level GO categories as well as the most represented GO-term are indicated. Four of those clusters are related to “response to stress”, and more precisely to “response to wounding”. Three other clusters are related to “immune system processes”. Among the remaining ones are found biological processes involved in “blood vessel development”, “regulation of plasma lipoprotein particle levels”, “cholesterol transport”, “locomotion”, “secretion”, “negative regulation of peptidase activity” or more general catabolic processes like “carbohydrate derivative catabolic processes” and “NAD metabolic process”.

**Figure 2.**
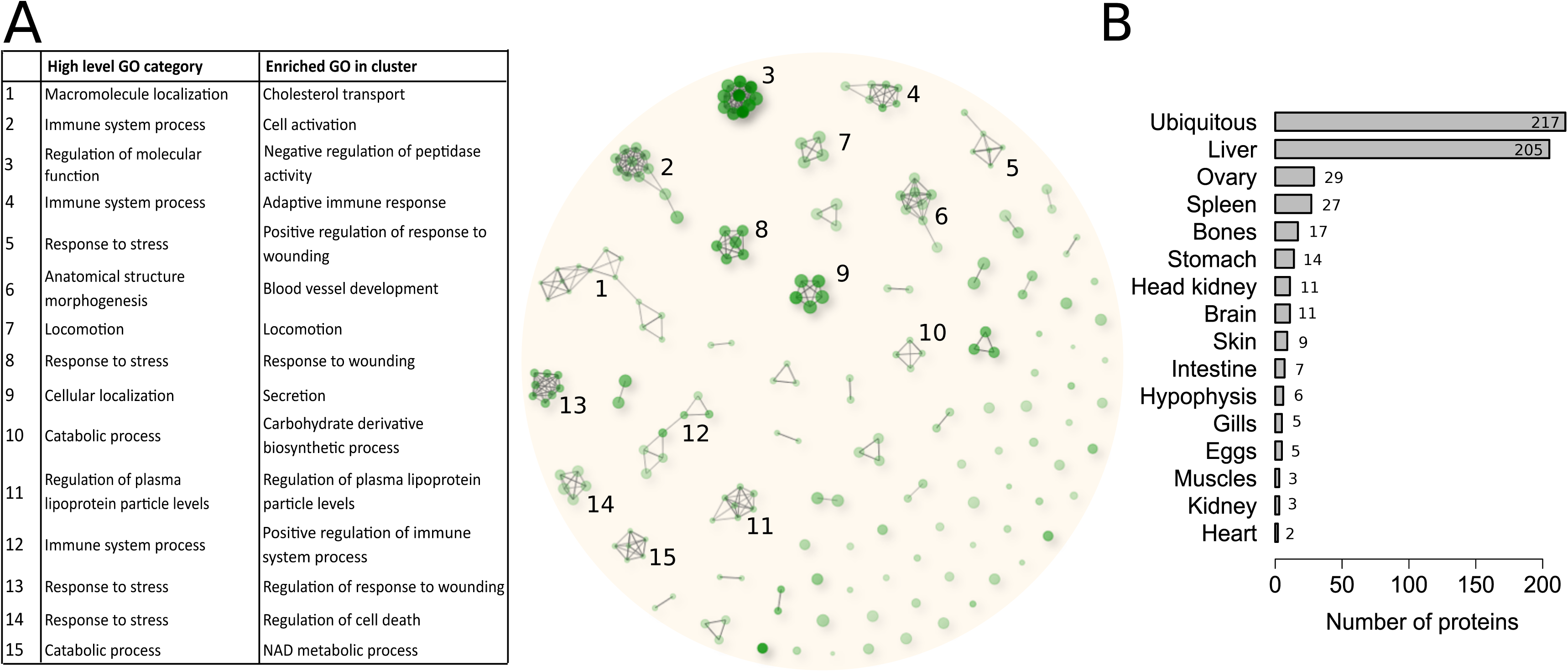
Proteomic characterization of rainbow trout coelomic fluid. Hierarchical clustering of the 500 most enriched biological processes GO-terms (FDR <0.05) in the rainbow trout CF proteome (A). Related GO-terms in a cluster share at least 70% of their proteins. Proteins of the main clusters (clusters with at least 4 GO-terms) were further subjected to GO enrichment analysis to identify high-level GO categories representing the different clusters, as well as the most represented biological processes in each cluster. Main tissue of expression of the rainbow trout CF proteins (B).

The amino acid sequences of all the proteins identified in the trout CF were blasted against the PhyloFish transcriptomic database ^29^. For each protein, the main tissue of expression of the corresponding contig was manually checked (Figure 2B, Table S3). Proteins were classified based on the tissue/organ exhibited predominant expression. Proteins exhibiting a ubiquitous expression were the most abundant. In addition, a high proportion of proteins exhibited a predominant expression in the liver. This observation is consistent with the previously reported presence of egg yolk proteins in rainbow trout fluid^24^. We also identified 29 proteins that appear to be predominantly expressed in the ovary. This is consistent with the ovarian origin of several CF proteins previously described in salmonids, including progastricsin^10^ and trout ovulatory proteins^8,9^.

### Identification of salmonid specific CF proteins

Extended egg viability is a salmonid-specific trait. In order to identify proteins conferring this specific egg conservation property to the salmonids CF, a proteomic comparison of three different salmonid species CF (Rainbow trout, *Oncorhynchus mykiss;* Atlantic salmon*, Salmo salar* and brown trout*, Salmo trutta*) to three non-salmonid teleost species OF (Goldfish, *Carassius auratus;* Common carp, *Cyprinus carpio;* and Pikeperch, *Sander lucioperca)* was performed. Together, 511 proteins were identified in Atlantic salmon CF, 388 in brown trout CF, 876 in goldfish OF, 423 in common carp OF and 392 in pikeperch OF (Table S1). To facilitate inter-species comparisons, all identified proteins were mapped to teleost common ancestors using the Genomicus^30^ database in order to identify groups of orthologous proteins. A total of 368 unique common ancestors were found in rainbow trout CF, 311 in Atlantic salmon CF, 299 in brown trout CF, 527 in goldfish OF, 420 in common carp OF and 295 in pikeperch OF (Table S1).

Quantitative proteomic comparison was performed using those common ancestors IDs. For statistical purposes, only common ancestors identified in at least 2 species of one of the 2 groups were kept: 303 for the salmonid group and 268 for the non-salmonid group. Each group shared roughly half of its identified common ancestors with the other group (Figure 3A). From the 419 common ancestors quantified, 78 were found significantly enriched in the salmonid group in comparison to the non-salmonid group (Figure 3B, Table S4). In rainbow trout CF, 97 proteins corresponding to 66 common ancestors were found. Proteins corresponding to the 12 remaining enriched common ancestors were only found in the CF of the two other salmonid species (Table S4).

**Figure 3.**
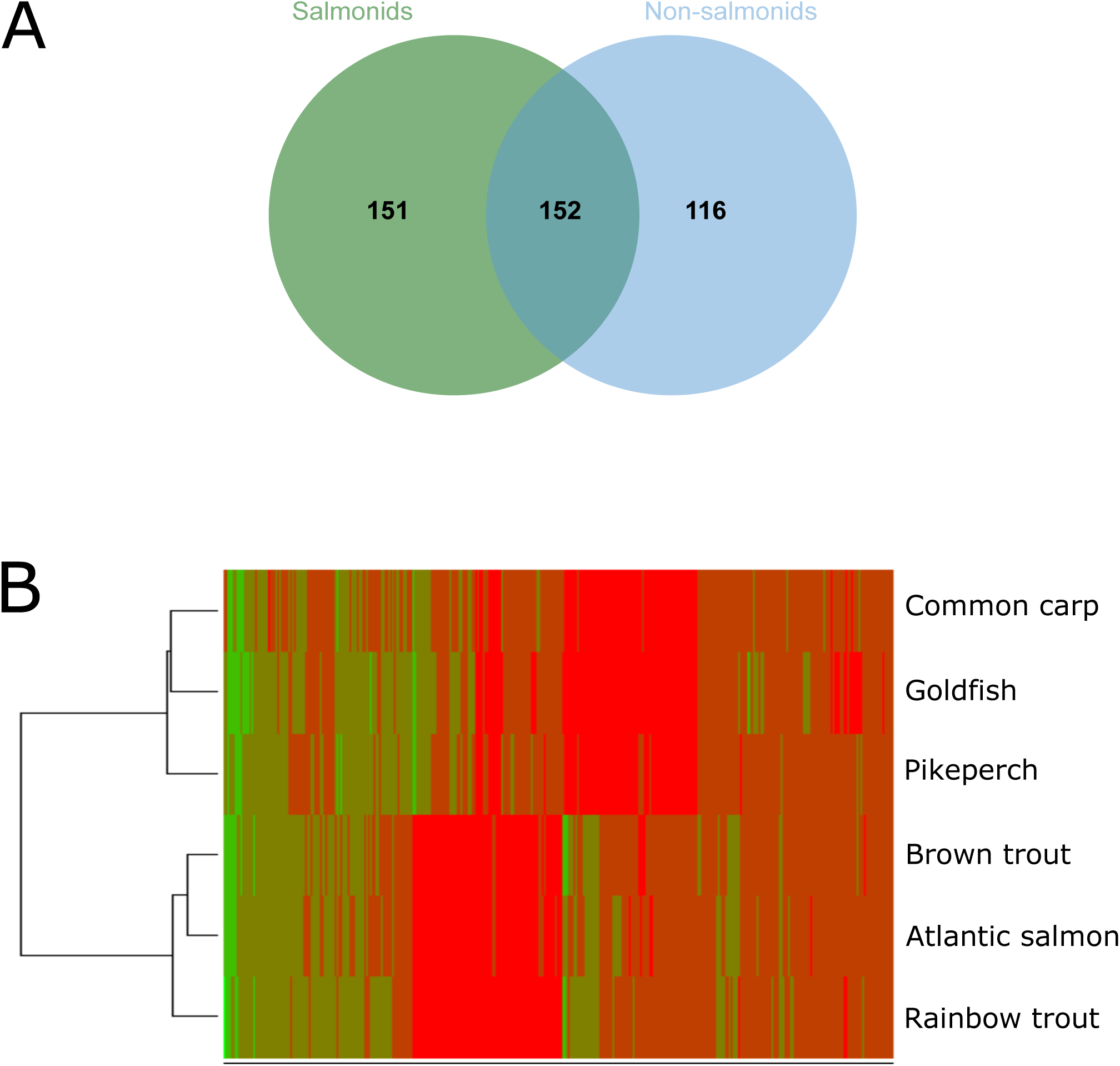
Differential proteomic comparison of salmonids CF and non-salmonids teleost OF. Venn diagram showing the overlapping between the different species CF and OF proteomic composition (A). Venn diagram showing the overlapping between the salmonids group CF and the non-salmonids teleost group OF composition (B). Venn diagram showing the overlapping between the 3 species CFs of the salmonids group (C). Venn diagram showing the overlapping between the 3 species OFs of the non-salmonids teleost group (D). Heat map and hierarchical clustering of the quantified common ancestors in the 6 species of the 2 groups (salmonids and non-salmonids teleost) (E). Red indicates high abundance while green indicates low abundance.

### Identification of rainbow trout CF proteins candidates involved in egg conservation

The proteomic comparison of salmonid CF to non-salmonid OF led to the identification of proteins specifically present in salmonids CF. This set of 78 common ancestor genes could be playing an important role in maintaining egg viability and ability to be fertilized after ovulation. In addition to this evolutionarily-supported analysis, we used a functional angle. We speculated that different protein fractions of the salmonid CF could play different role in preserving egg viability and ability to be fertilized. In order to further identity proteins involved in maintaining egg viability, rainbow trout CF was analytically fractionated. Using gel filtration column, CF proteins were separated in 30 different fractions according to their size. Then, the ability of the different fractions to preserve trout egg quality after 3 days of storage was evaluated. Storage in non-fractionated CF was used as a positive control.

Strong differences in embryonic survival rate could be observed between the fractions (Figure 4A), confirming our working hypothesis that different CF fractions had different contribution to CF biological activity. Three main peaks of egg survival rate were observed in fractions F3, F12 and F28 with 42%, 61% and 61.5 % of eggs reaching eyed-stage, respectively. These percentages were not significantly different from the one obtained with the non-fractionated CF. Proteins present in those fractions were subsequently analyzed by mass spectrometry. To tentatively identify candidate proteins involved in egg preservation, the two groups of fractions with the highest egg developmental rates, F12 (group F11-F13) and F28 (group F27-F29), were selected for further analysis.

**Figure 4.**
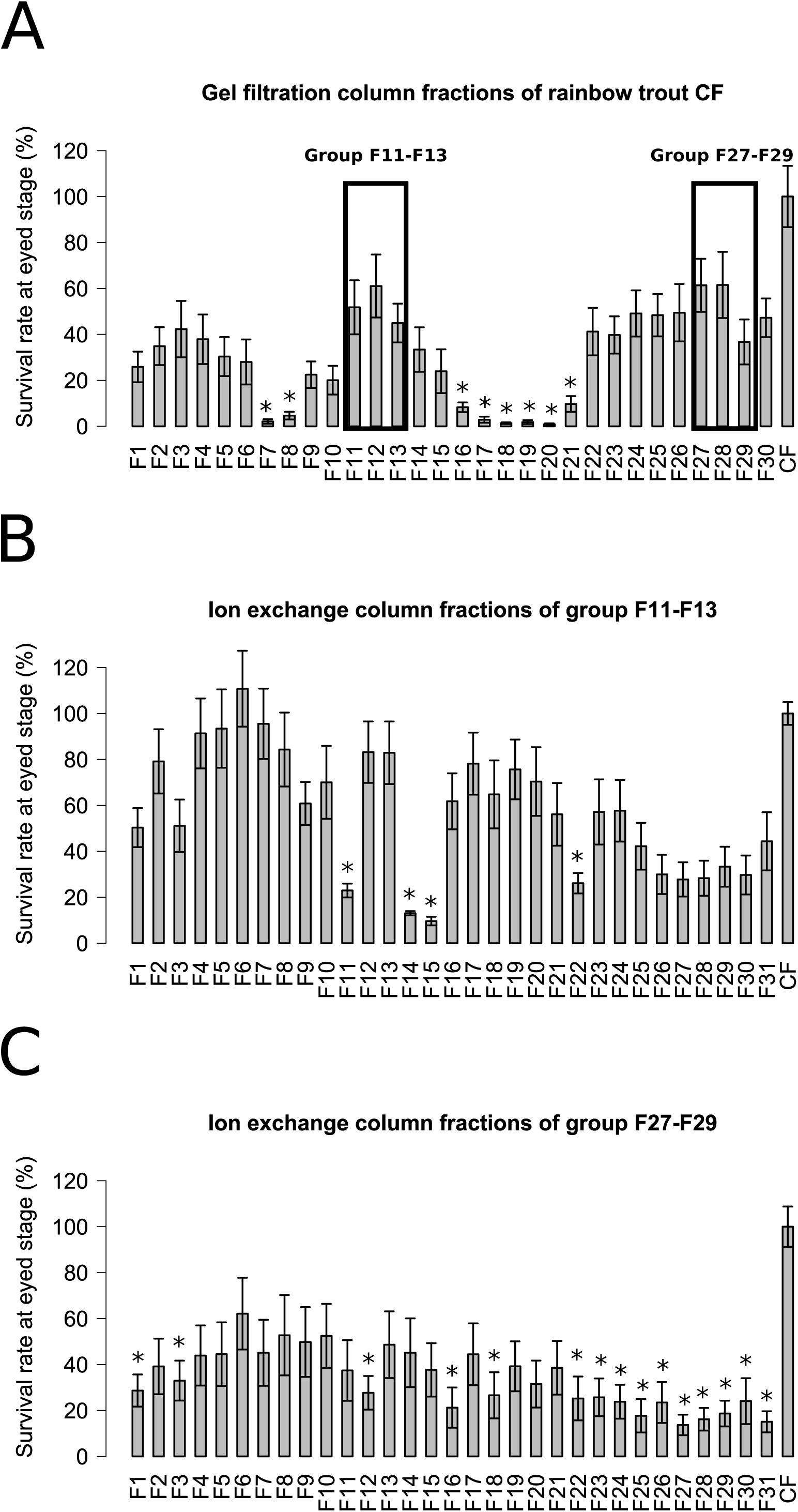
Using gel filtration, coelomic fluid proteins were separated into 30 different fractions according to their size (A). Proteins from the group F11-F13 were separated according to their charge onto an anion exchange column (B). Proteins from the group F27-F29 were separated according to their charge onto an anion exchange column (C). Percentage of eggs reaching eyed stage after 3 days storage in the different fractions were evaluated. All bars represent the means ± sem of experiments conducted on the eggs of 4 females. *Significantly different from CF, t-test (p-value<0.05).

For these two fractions, a second fractionation was performed. An ion exchange column was used to separate the proteins according to their charge. The egg viability conservation property of the different fractions obtained was evaluated as previously described. The results for the group F11-F13 fractionation are presented in figure 4B. Three main pics of egg developmental rate were observed in the fractions F6, F12 and F17 with 111%, 83% and 78% of eggs reaching eyed-stage respectively. The percentages of these 3 main peaks were not significantly different from the percentage obtained with the non-fractionated CF. The results for the group F27-F29 fractionation are presented in figure 4C. Observed survival rates were lower than in the two previous fractionation experiments. However, a pic of egg developmental rate, reaching 62%, was observed in the fraction F6. This egg developmental rate was not significantly different from the non-fractionated trout CF. Proteins present in the fractions allowing good egg developmental rate were identified by MS: F6, F12 and F17 for the group F11-F13, and F6 for the group F27-F29. As a comparison, fractions with egg developmental rate significantly lower than with non-fractionated CF were also analyzed by MS. Those fractions were selected preferentially “close” to the fractions allowing good egg conservation during the fractionation process. Fractions F11, F14 and F22 were selected for comparison with the F11-13 group while fraction F12 was selected for comparison with the F27-F29 group. Normalized spectral counts (N-SC) ratios of good versus bad egg conservation fractions were then calculated. A total number of 376 proteins were found specifically present or highly enriched (N-SC ratios > 2) in the fractions allowing good egg viability conservation (Table S5).

Interestingly, 50 of those proteins specifically present or enriched in the rainbow trout CF fractions allowing egg viability conservation were also found among the salmonid specific CF proteins (Table S5). They constitute a list of potential candidates conferring the exceptional egg conservation property to salmonid CF.

### Functional analysis of the N-acetylneuraminic acid synthase a (Nansa) protein

Among the potential candidates identified, some are known to play a role in the ovulation process. We decided to focus on a specific protein not previously reported in salmonids CF, and with no known function in the ovulation process: the sialic acid synthase protein (XP_021429200.1). According to Ensembl, this protein is encoded by the N-acetylneuraminic acid synthase a gene (*nansa*) (ENSOMYG00000017265) that is orthologous to the human *NANS* (ENSG00000095380) gene. Sialic acid synthases are responsible for the formation of sugars on glycoproteins in eukaryotic cells and are involved in fertilization, embryogenesis, and differentiation ^31–33^. The identified Nansa protein has a NeuB domain and a SAF NeuB-like domain. NeuB is the catalytic domain involved in N-acetylneuraminic acid (Neu5Ac) in procaryotes, whereas SAF NeuB-like domain would allow the binding to substrates. SAF NeuB-like domain is part of the superfamily domain SAF which also includes the domains similar to fish antifreeze proteins according to the NCBI conserved domain database^34^.

According to Phylofish database^29^, the contig corresponding to Nansa is mainly expressed in the ovary (Table S3). Using quantitative RT-PCR we observed that *nansa* was strongly and predominantly expressed in the ovary, but also in the testis (Figure 5A). In the ovary, *nansa* is progressively down-regulated (Figure 5B) following ovulation. Nansa protein profile observed in ovary and in CF are consistent with those gene expression results. Our western blot analysis showed that Nansa is highly expressed in the ovary and the CF at ovulation. Nansa protein abundance then progressively decreased following ovulation (Figure 5C and D).

**Figure 5.**
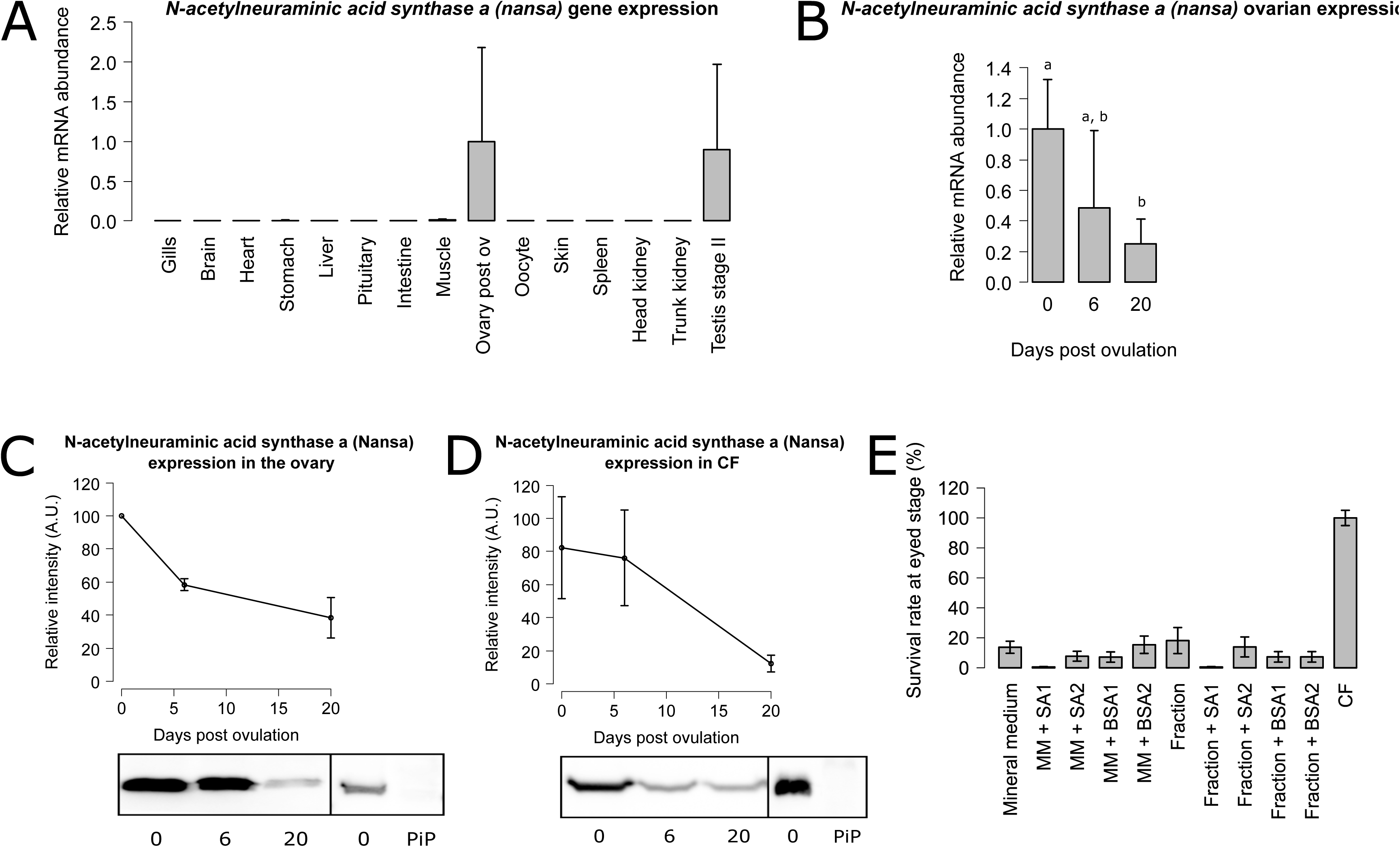
A) Tissue expression profile of *N-acetylneuraminic acid synthase a* (*nansa*) in rainbow trout. Tissues were sampled from 3 different fish. The *nansa* mRNA abundance was determined by qPCR and normalized to the abundance of 18S. Abundance was set to 1 for the maximal gene expression and data are expressed as percentage of transcript abundance. B) Ovarian expression profile of *N-acetylneuraminic acid synthase a* (*nansa*) gene in the ovary of rainbow trout on the day of ovulation and 6 and 20 days after ovulation. The *nansa* mRNA abundance was determined by qPCR and normalized to the abundance of 18S. Abundance was set to 1 for gene expression on the day of ovulation and data are expressed as percentage of transcript abundance on this day. For each ovary stage, ovaries were sampled from 3 separate females. Different letters indicate significant differences (t-test, p<0.05). C) Representative western blot analysis with antibody against N-acetylneuraminic acid synthase a (Nansa). The ovary samples were analyzed at 0, 6, and 20 days after ovulation. Band intensity was normalized by setting the maximal signal obtained to 100%. (Upper panel). The arrow indicates the N-acetylneuraminic acid synthase a (Nansa) band at 40kDa. The positions of the molecular weight markers are indicated on the left. Pre-immune serum (PiP) was used as a negative control (Low panel). D) Representative western blot analysis with antibody against sialic acid synthase. CF samples were analyzed at d0, d6 and d20 post ovulation. Band intensity was normalized by setting the maximal signal obtained to 100% (Upper panel). The arrow indicates the sialic acid band at 40kDa. The positions of the molecular weight markers are indicated on the left. Pre-immune serum for antibody sialic acid synthase-like was represented (PiP) (Low panel). E) Percentage of rainbow trout eggs reaching eyed stage after 3 days of *in vitro* storage in mineral medium, mineral medium complemented with 2 different concentrations of N-acetylneuraminic acid synthase a (Nansa) (SA1, 4 µg/mL; SA2 40 µg/mL), in a CF fraction not allowing good egg conservation, the same fraction complemented with 2 different concentrations of N-acetylneuraminic acid synthase a (Nansa) (SA1, 4 µg/mL; SA2 40 µg/mL), and in CF before fertilization. BSA was used as a negative control for mineral medium and fraction complementation at the same concentrations as sialic acid synthase-like (BSA1, 4 µg/mL; BSA2 40 µg/mL). All bars represent the means ± sem of experiments conducted on the eggs of 3 females.

To evaluate the contribution of sialic acid-synthase protein to egg conservation, rainbow trout eggs were stored for 3 days at 12°C in the dark, in MM complemented with two different concentrations of sialic acid-synthase or in a CF fraction not allowing a good egg conservation complemented with two different concentrations of sialic acid-synthase, before fertilization. As a control, eggs were also incubated in MM or in CF fraction complemented with BSA. Complementation with sialic acid-synthase did not improve embryonic survival rate (Figure 5E).

## Discussion

### Proteomic characterization of the rainbow trout CF

In our study, we demonstrate that using mineral medium mimicking the chemical composition of rainbow trout CF for egg storage is not sufficient to preserve egg viability and ability to be fertilized. We showed that organic compounds, and more specifically proteins, confer egg conservation properties to rainbow trout CF (Figure 1). To better understand this biological property, the rainbow trout CF proteomic composition was analyzed. This led to the identification of 573 proteins (Table S1), which is consistent with data recently obtained for Chinook salmon CF proteome^16^. However, compared to the previously 54 proteins identified in the rainbow trout CF ^26^, the present analysis provides a more comprehensive characterization of the rainbow trout CF proteome. Together, these results indicate that CF is a complex biological fluid.

Gene ontology analysis of the rainbow trout CF proteome revealed fifteen clusters of biological processes that are highly consistent with previous proteomic analyses of rainbow trout and chinook salmon ^26,27^ (Figure 2, Table S2). Four GO clusters were related to “response to stress” and more precisely to “response to wounding”. This group of proteins are involved in response to injuries, which is consistent with the response to the follicle epithelium rupture at ovulation. Three other clusters were related to “immune system processes”. In this group are notably found proteins of the Serpin family, Fibrinogen proteins, proteins of the complement or the von Willebrand factor. This group of proteins would be directly involved in egg conservation and more precisely in the egg protection against infections. Indeed, it was previously demonstrated that the trout CF has anti-bacterial property ^9^. However, complement proteins, which are present in this group, are not only restricted to immunity. They are also frequently associated to ovarian function. C3 has been identified as a potential biomarker for in vitro fertilization (IVF) outcome^35^ while iC3b, its product of cleavage, is involved in oocyte maturation in pigs^36^. Several complement proteins have been proposed to be oocyte maturation factors in human^37^. All the above-listed proteins are acute phase proteins (APP) that are expressed in response to injuries or infections. In mammals, the ovulation process shares similarities with the inflammation process, and numerous APP are commonly involved in both processes^38,39^. Similarly to mammals, it has been suggested that ovulation in fish would also be an inflammation-like process ^40,41^. Together, these observations are consistent with the identification of APPs in the two main represented biological processes in the rainbow trout CF : “response to wounding” and “immune system processes”.

Two GO clusters were related to lipids binding and transport: “cholesterol transport” and “regulation of plasma lipoprotein particles level”. Lipids play important dual functions during oogenesis. Cholesterol is an essential substrate for normal steroidogenesis and consequently for normal follicle development and ovulation. Furthermore, lipids are sequestered from the VLDL plasma and incorporated in the growing oocyte in the form of lipid granule where they constitute an energy reserve for the future developing embryo. As in previous trout and salmon CF proteomic studies, the most abundant proteins that we identified were vitellogenins (Table S1). Those glycolipophosphoproteins are the major egg yolk proteins. After incorporation in the growing oocyte, they are further processed and also constitute a metabolic energy reserve for the future growing embryo.

Among the remaining GO cluster are notably found “blood vessel development” and “regulation of peptidase activity” processes. Blood vessel development is important for the local delivery of proteins involved in the ovulation process or necessary to the growing oocyte. In the GO “regulation of peptidase activity” are found proteases and their inhibitors involved in the ECM remodeling and degradation like MMP9 and TIMP2. This process is necessary to the follicular rupture. In this GO term are also found proteases involved in the coagulation process, in response to the follicle rupture.

### Identification of candidate CF proteins involved in egg conservation

More than a comprehensive proteomic characterization of the rainbow trout CF, the main goal of this study was to identify proteins conferring its exceptional egg preservation ability to salmonid CF. We first used an evolutionary approach to identify proteins specifically enriched in salmonid CF. To our knowledge, this is the first study comparing the proteomic composition of CF/OF in different fish species. The main challenge was to find a way to perform inter species comparisons. To address this issue, all identified proteins were mapped to teleost common ancestors using the Genomicus database that provides synteny-supported orthology assignments. Then, species were separated in 2 groups: salmonid species and non-salmonid species. This comparison led to the identification of 78 unique teleost common ancestors significantly enriched in the CF of salmonid species (Figure 3, Table S4) in comparison to non-salmonid species.

We then used a functional approach to identify CF proteins specifically involved in egg conservation using a biological test. This analysis was conducted in rainbow trout. For this approach, rainbow trout CF was fractionated to allow an independent evaluation of the biological activity of the different fraction. This led to the identification of 376 rainbow trout CF proteins specifically enriched in fractions allowing good egg conservation (Figure 4, Table S5).

After compiling the results from evolutionary and functional analyses, 50 rainbow trout CF proteins were commonly identified by both approaches (Table S5). We speculated that these proteins are major players of egg viability preservation. Based on their functions, they can be separated in different categories that will be discussed below.

#### Immunity

This category includes complement system complement C1q-like (C1ql1), complement factor D (Cfd), complement factor H (Cfh), and precerebellin-like protein precursor (Cbln1). Cbln1 has previously been associated with acute phase response in rainbow trout^42^. Together with previous proteomic analyses ^16,26^ our data show that proteins involved in immunity are abundant in the CF of salmonid fish. In addition, we provide functional evidence that these players would play important roles in preserving eggs viability when eggs are held in CF.

#### Calcium-dependent and calcium-binding proteins

Our results also suggest that several calcium-dependent and calcium-binding CF proteins play important roles in preserving egg viability, including 45 kDa calcium-binding protein, calcium/calmodulin-dependent protein kinase type II subunit delta (Camk2d), calumenin-A (Calu), ependymin ^43^, lithostathine-1-like^44^, and B-cadherin. Calcium (Ca2+) is an important parameter modulating sperm velocity. Indeed, there is a negative correlation between duration of sperm motility and OF Ca2+ concentration ^21^. Chemical composition, including Ca2+ concentration, of several salmonids CF has been extensively studied ^20,21^. In fact, a high proportion of calcium would not be free but bound to proteins ^21^. The calcium-binding protein that we identified in our study might be involved in the regulation of free Ca2+ concentration in salmonid CF. High levels of calcium binding proteins in CF would presumably help extending sperm motility and increase fertilization success. Consistent with this hypothesis is the abundance of several calcium-binding proteins in Chinook salmon CF^16^.

#### Sialic acid related proteins

Several sialic acid related proteins were identified in our analysis, including sialic acid synthase, alpha-N-acetylgalactosaminide alpha-2,6-sialyltransferase 1-like. Sialic acids are 9 carbons sugar structures present as terminal residues of glycoproteins. Specific trout egg glycoproteins are progressively sialylated during oogenesis^45^. Several sialyltransferases are involved in this process, notably alpha-N-acetylgalactosaminide alpha-2,6-sialyltransferase 1. It was hypothesized that sialic acid structures could have several functions in egg protection including protection from maternal complement attack^33^. Interestingly, complement factor H, that we also identified in our screen, is involved protection from maternal complement. Indeed, factor H down-regulates activation of the alternative complement pathway. By binding sialic acids expressed at cell surface, it allows self-recognition and prevent cells to be attacked by the complement system^46^. Sialic acid might also be involved in calcium binding. Ependymin, one of our candidates, is a Ca binding protein that is a sialylated. It seems that sialic acid residues are responsible for this Ca binding property ^43^.

#### Lipid transport and catabolism

Our screen also led to the identification of several proteins involved in lipid transport and catabolism/hydrolysis including apolipoprotein B-100, beta-2-glycoprotein 1, carboxylesterase 3, phospholipase B1, and prosaposin. As previously discussed, lipids play important function during the oogenesis. Apo B-100 is involved in lipid transport, and notably cholesterol, the substrate for steroid hormones synthesis by the ovarian follicle^41^. It may also be involved in the storage of neutral lipid in the form of lipid globules in the oocyte cytoplasm. This step would also require the action of lipid hydrolases^47,48^. The identification of proteins of this category in fish ovarian fluid is therefore not surprising. However, it is not clear why they are specifically enriched in Salmonids CF and in CF fractions allowing a better egg conservation. Their role in egg protection is currently unknown and would require further investigations.

#### Extracellular matrix (ECM) proteins and ECM degrading enzymes

Many ECM proteins and ECM degrading enzyme were also identified as possible major players responsible for salmonid CF ability to preserve egg viability including cathepsin K, matrix metalloproteinase-9 (Mmp9), olfactomedin-4, versican b, thrombospondin-1, cytoskeleton proteins (tubulin alpha chain, scinderin like b, plastin-2) and cell adhesion proteins (integrin beta-1, B-cadherin). These results are consistent with the previous report of protease activity and notably collagenase activity in salmonid CF^20^. Mmp9 is a gelatinase expressed in the teleost ovary^49^ and involved in the ovulation process^50^. CtsK is a collagenase involved in bone resorption^51^. CtsK has been detected in the mouse ovary but its involvement in ECM degradation in the ovary has not been demonstrated ^52^. At the time of ovulation, proteases play a role in the lysis of the follicular epithelium to allow the release of mature oocytes into the coelomic cavity. Proteolytic enzymes and components of the extracellular matrix might also be released in the coelomic fluid during this process. Whereas their abundance in salmonids CF can be explained, their role in the egg conservation process remains unclear. However, some of those proteins are associated to oocyte quality or fertilization process in vertebrates. In humans, MMP-9 levels in follicular fluid are positively correlated with oocyte quality and fertilization rate^53^. Furthermore, spermatozoa capacitation appears to require interaction with ECM proteins^54^. In summary, while their precise contribution is unclear, it cannot be excluded that some of the identified ECM candidate proteins might contribute to preserving unfertilized egg viability.

### Preservation of egg viability and quality is a multi-protein process

Our study indicates that protective role of CF over the egg viability during the *in vivo* post-ovulatory storage is associated with the presence of multiple abundant proteins. Several studies have previously led to the identification of abundant CF proteins, including CF-specific proteins^10,55,56^. In the present study we have specifically investigated sialic acid synthase (Nansa) due to its presence in a CF fraction exhibiting high biological activity, because several sialic acid related proteins have been identified as being abundant only in salmonid species. Yet, supplementation of mineral medium with Nansa at two different concentrations did not improve egg viability and quality. While this could be due to methodological limitations, including concentration and biological activity, we can speculate that Nansa, alone, is not sufficient to preserve egg viability and quality. Together with the high number and functional diversity of proteins found in biologically active fractions and in the 3 salmonid species investigated these observations suggest that salmonid CF is a complex biological fluid. We propose that a significant number of proteins present in CF are collectively responsible for providing it unique ability to preserve egg viability and quality. Among these key players are proteins involved in immunity, calcium binding, lipid metabolism, proteolysis, extracellular matrix and sialic acid metabolic pathway.

## Methods

### Coelomic fluid, egg and sperm collection

Two-year old rainbow trout (*Oncorhynchus mykiss*) were obtained from the INRAE PEIMA experimental fish farm (Sizun, France) approximately 1 month before spawning. Fish were held during spawning season in a recirculated water system at 12°C under a natural photoperiod (ISC LPGP, Rennes, France) until ovulation. Twice a week, fish were anaesthetized using tricaine methanesulfonate (MS222) at a concentration of 80 mg/L and checked for ovulation using gentle abdominal pressure. At the time of detected ovulation, eggs and coelomic fluid were collected by hand stripping (*i.e*., applying pressure onto the abdomen). Eggs were separated from CF with using a mesh screen. A turbidity test was performed as previously described ^18^ for each collected CF to avoid any bias due to damaged or leaking eggs and only non-precipitating CF were kept. CF sample were then centrifuged at 3000g for 15 min at 4°C to remove red blood cells and debris (pure CF). The collected supernatants were stored at -20°C until use. When mentioned, the supernatants were either boiled at 95°C for 10 min to denature proteins (boiled CF) or ultracentrifuged 1h, at 100 000 g, at 4°C to float up lipids (CF ultracentrifuged). CF depleted in lipids was collected from the bottom of the tube with the help of a needle and stored at -20°C until use. Before storage at -20°C, protein concentration of the CF was determined using the Coomassie (Bradford) Protein assay Kit (Thermo Fisher Scientific, Rockford, IL, USA) and bovine serum albumin (BSA) as a calibration standard. To collect sprm, the genital region was wiped and sperm was obtained by manual stripping. A pool of sperm was made from 3 males and stored on ice until fertilization. Atlantic salmon (*Salmo salar*) CF and Brown trout (*Salmo trutta*) CF were obtained as described above for rainbow trout. Goldfish (*Carassius auratus*) were obtained from the INRAE U3E facility in Rennes, France. Two-year-old fish raised in outdoor tanks were transferred into 0.3 m^3^ tanks and reared several weeks in recycled water at 14°C under a 14 h light and 10 h dark photoperiod (ISC-LPGP, Rennes, France). Fish were fed with carp pellets at a ratio of 1% of body weight. Gamete release and ovarian fluid were obtained according to *Depince et al* ^18^. Common carp (*Cyprinus carpio*) OF samples were provided by the University of South Bohemia in České Budějovice, Faculty of Fisheries and Protection of Waters, České Budějovice, Czechia. Pikeperch (*Sander lucioperca*) OF samples obtained from domesticated pikeperch broodstock reproduced following standardized controlled reproduction protocol as previously described^57^. Briefly, fish were held at the Asialor fish farm where they were subjected to a specific annual photo-thermal program enabling proper course of gonadal development. Just prior to spawning ovulation was stimulated using salmon gonadoliberine analog (50 µg/kg of body weight; Bachem, Switzerland). When ovulation was observed, eggs were hand-stripped into a dry container and OF was aspirated with a pipet into cryotubes which were immediately after snap frozen in liquid nitrogen. OF samples were then stored at -80°C. For the current experiment OF was collected only from egg portions characterized by highest quality (fertilization rate above 70%). CF and OF samples collected from those different species were processed as described above for rainbow trout CF.

### Tissue collection

For tissue collection, female rainbow trout were euthanized using a lethal dose of tricaine methanesulfonate (MS222) at a concentration of 200 mg/L. Ovaries were dissected and ovarian samples were either frozen in liquid nitrogen and stored at -80°C until RNA extraction or homogenized in RIPA buffer (NaCl 150 mM, deoxycholic acid 0.5%, NP40 1%, SDS 0.1%, Tris 50 mM, pH 8) using a Precellys Evolution Homogenizer (Ozyme, Bertin Technologies) to extract proteins. After centrifugation, protein concentration in the supernatant was determined using the Coomassie (Bradford) Protein assay Kit (Thermo Fisher Scientific, Rockford, IL, USA) and bovine serum albumin (BSA) as a calibration standard. Supernatants were then stored at -20°C until use.

### Biological test to assess egg viability preservation potential

*In vitro* eggs storage was performed, under sterile conditions in 6-well polystyrene culture plates (Falcon). Before storage, eggs were washed with filter-sterilized (0.22 µm) trout mineral medium (MM) ^28^ (124.1 mM NaCl, 5.1 mM KCl, 1 mM MgSO4.7H_2_O, 1.6 mM CaCl_2_.2H_2_O, 5.6 mM Glucose, 26.5 mM Hepes, pH 8) to remove residual CF. A volume of 3 mL of medium (CF, boiled CF, ultracentrifuged CF, diluted CF, CF fractions, or MM) was used in each well and eggs were added at a ratio of 35 eggs per well. Technical duplicates (2 × 35 eggs) were used for each condition. Plates were then stored 3 days in the dark, in a 12°C incubator, under gentle and constant agitation (50 rpm) until fertilization. For CF fractions complemented with BSA or sialic acid synthase (Nansa), two concentrations were tested: 4 or 40 µg/mL. The sialic acid synthase protein (NCBI accession # XP_021429200) was commercially produced using the insect expression vector pFastBac1 (GenScript U488UFB180).

### Fertilization and developmental success monitoring

Fertilization was performed as previously described with minor modification ^7^. Eggs were transferred into plastic glass tubes. Sperm was added (5µL) along with Actifish (IMV, L’aigle. France). Tubes were gently swirled. After 5 min, Actifish was discarded, eggs were transferred into plastic incubators and held in the dark at 10°C in a recirculated water system during early development. Total number of eggs was counted. The number of embryos reaching eyed stage was counted 29 days after fertilization (dpf).

### RNA extraction and reverse transcription

RNA extraction was performed as previously described with minor modifications ^58^. Briefly, tissues were homogenized in TRI Reagent (TR118, Euromedex) at a ratio of 100 mg per mL of TRI Reagent. Total RNA was then extracted using the TRI Reagent procedure and resuspended in water. Reverse transcription was performed from 1.6 µg of total RNA using Maxima First strand kit (ThermoScientific) according to manufacturer’s instructions and using the following steps: 10 min at 25°C, 30 min at 60°C, and 5 min at 85°C. Control reactions were run without reverse transcriptase and used as negative real-time PCR controls.

### Real-time PCR

RT products and negative control reaction products were diluted to 1/25 (1/2000 in the case of the *18S* gene) and 4 µL were used in each PCR reaction. For each sample, triplicates were analyzed. Real-time PCR was performed using a real-time PCR kit provided with SYBR green fluorophore (Powerup SYBER Green Master Mix, Applied Biosystems) with 600 mM of each primer. For sialic acid synthase, the following primers were used ACCTGGCCCATCACTTTACG/ACATCACCCAGTGCCATGTT. Relative abundance of target cDNA in the sample was evaluated from a serially diluted cDNA pool (standard curve) using the StepOne+ cycler software (Applied Biosystems). Real-time PCR data were all normalized to *18S* RNA abundance in the samples.

### SDS-PAGE and Western Blot

SDS-PAGE and western blot were performed as previously described ^58^ with minor changes. Protein samples were diluted in Laemmli buffer ^59^ at a final concentration of 1 µg/µL and for each sample, 20 µL were loaded per well. A custom-made antibody directed against peptide CKVGEPRGVSPEDMG of the sialic acid synthase (NCBI accession # XP_021429200) was purchased (GenScript U881EER070) and used at a dilution of 1/2000. Anti-rabbit horseradish peroxidase (HRP)-conjugated antibody (Jackon Immunoresearch, West Grove, USA) was used as secondary antibody at 1/5000 dilution. Detection was performed with Uptima Uptilight chemiluminescent revelent kit (Uptima-Interchim) using a Fusion FX7 imager (Vilbert Lourmat) with the Fusion software (v 15.11). Band intensities were measured using the ImageJ software. For each temporal profile, intensities were normalized by setting the maximal signal obtained to 100%.

### Sample preparation of rainbow trout coelomic fluid before nanoLC-MS/MS analysis

#### HPLC fractionation of the rainbow trout coelomic fluid

Coelomic fluid (CF) samples were desalted and concentrated by diafiltration (10kDa Vivaspin, Sartorius) according to manufacturer instructions and then fractionated by UHPLC using sequential chromatography (gel filtration and anion exchange). First, samples (60 mg of proteins) were resuspended in 2 mL of 10 mM sodium phosphate / 0.15 mM sodium chloride, pH 6.8 and loaded onto a gel filtration column (TSK gel G3000SW 21.5 mm i.d.x30cm, 13 µm; Tosoh Bioscience, Tokyo, Japan) at a flow rate of 4 mL/min. Bound proteins were eluted isocratically using the phosphate buffer during 40 min. We monitored the UV absorbance of the eluent at 210 and 280 nm. Thirty fractions were collected and systematically assessed for egg quality preservation with in vitro egg storage test followed by fertilization and developmental success monitoring.

Fractions of interest were further selected for a second chromatography fractionation. They were desalted and resuspended in 250µL of 10mM Tris pH 7.6. The samples were then loaded onto an anion exchange column (ProSwift SAX-1S 4.6×50 mm PK/SS; Thermo Fisher Scientific, Courtaboeuf, France) at a flow rate of 1 mL/min. The mobile phases used were (A) 10 mM Tris, pH 7.6 and (B) 1M NaCl in 10mM Tris, pH 7.6. After sample loading, the gradient applied was (min:%B); 0:0, 5:2, 30:50, 40:100, 50:100, 60:0, 120:0. Fractions were collected every 60 s from 0 min to 46 min of the gradient, resulting in 46 fractions. The detection of protein was carried out at 210 nm and 280 nm. These fractions were also tested on ovulated eggs and proteins in fractions of interest were further analyzed by mass spectrometry after liquid trypsin digestion for protein identification.

The fractions of interest were desalted by diafiltration (10 kDa Vivaspin), and resuspended in 20 µL of 50 mM ammonium bicarbonate/0.01% ProteaseMax (Promega, Charbonnières-les-Bains, France) then incubated in 2.5 µL of 65 mM DTT at 37°C for 15 min followed by incubation at room temperature in the dark for 15 min after addition of 2.5 µL of 135 mM iodoacetamide. Finally, 2 µL of sequencing grade modified trypsin (Promega) at a concentration of 0.1 µg/μL and 23 µL of 50mM ammonium bicarbonate/0.01% ProteaseMax were added. The trypsin digestion was performed overnight at 37°C and resulted peptides were analyzed by nanoLC-MS/MS using a short nanoLC gradient.

#### Proteomic comparison of salmonid CF to non-salmonid OF

Samples of salmonid CF and non-salmonid OF (2 µg of proteins for each sample) were subjected to a short migration on a NuPAGE system (Invitrogen, 3 minutes at 200 V-400 mA - 23 W) according to the manufacturer’s instructions, in order to stack the proteins of each sample in one band. After coloration using the EZBlue Gel Staining Reagent (Sigma-Aldrich, Saint-Quentin Fallavier, France) bands were excised from the gel and processed for tryptic digestion and peptide extraction as previously described ^60^. Resulting peptides were analyzed by nanoLC-MS/MS using a short nanoLC gradient (see below).

#### NanoLC-MS/MS analysis

Trypsic peptide mixtures were separated onto a 75μm×250 mm IonOpticks Aurora 2 C18 column (Ion Opticks Pty Ltd., Bundoora, Australia). A gradient of basic reversed-phase buffers (Buffer A: 0.1% formic acid, 98% H2O MilliQ, 2% acetonitrile; Buffer B: 0.1% formic acid, 100% acetonitrile) was run on a NanoElute HPLC System (Bruker Daltonik GmbH, Bremen, Germany) at a flow rate of 300 nL/min at 50°C for the HPLC fractions and 400nL/min at 50°C for non-fractionated samples. For the HPLC fractions, the liquid chromatography (LC) run lasted for 40 min (2% to 11% of buffer B during 19 min; up to 16% at 26 min; up to 25% at 30 min; up to 85% at 33 min and finally 85% for 7 min to wash the column). For the non-fractionated samples, the liquid chromatography (LC) run lasted for 120 min (2% to 15% of buffer B during 60 min; up to 25% at 90 min; up to 37% at 100 min; up to 95% at 110 min and finally 95% for 10 min to wash the column).

The column was coupled online to a tims TOF Pro (Bruker Daltonik GmbH, Bremen, Germany) with a Captive Spray ion source (Bruker Daltonik). The temperature of the ion transfer capillary was set at 180^11^C. Ions were accumulated for 114 ms, and mobility separation was achieved by ramping the entrance potential from−160 V to−20 V within 114 ms. The acquisition of the MS and MS/MS mass spectra was done with average resolutions of 60,000 and 50,000 full width at half maximum (mass range 100–1700 m/z), respectively. To enable the PASEF method ^62^, precursor m/z and mobility information was first derived from full scan TIMS-MS experiments (with a mass range of m/z 100–1700). The quadrupole isolation width was set to 2 and 3 Th and, for fragmentation, the collision energies varied between 31 and 52 eV depending on the precursor mass and charge. Tims, MS operation and PASEF were controlled and synchronized using the control instrument software OtofControl 5.1 (Bruker Daltonik). LC-MS/MS data were acquired using the PASEF method with a total cycle time of 1.31 s, including 1 tims MS scan and 10 PASEF MS/MA scans. The 10 PASEF scans (100 ms each) containing, on average, 12 MS/MS scans per PASEF scan. Ion mobility-resolved mass spectra, nested ion mobility vs. m/z distributions, as well as summed fragment ion intensities were extracted from the raw data file with DataAnalysis 5.1 (Bruker Daltonik).

Generated spectra were analyzed with the Mascot database search engine (v2.6.2; http://www.matrixscience.com) for peptide and protein identification, using its automatic decoy database search to estimate a false discovery rate (FDR) and calculate the threshold at which the e-values of the identified peptides were valid. According to the samples, the spectra were queried simultaneously in their species corresponding proteome database (NCBI *Oncorhynchus mykiss* – 20200923 *-* 97743 sequences; NCBI *Carassius auratus* – 20180809 *-* 96703 sequences; NCBI *Salmo salar* – 20210421 *-* 97555 sequences; NCBI *Cyprinus carpio* – 20210512 *-* 63928 sequences; NCBI *Salmo trutta* – 20210424 *-* 87841 sequences; NCBI *Sander lucioperca* – 20200814 – 56557 sequences) and in a database containing known contaminants such as human keratins or porcine trypsin (247 sequences). Mass tolerance for MS and MS/MS was set at 15 ppm and 0.05lilDa, respectively. The enzyme selectivity was set to full trypsin with one mis-cleavage allowed. Protein modifications were fixed carbamidomethylation of cysteines, variable oxidation of methionine. Identified proteins are validated with an FDR < 1% at PSM level and an e-value < 0.01, using Proline Studio v2.1.2 software ^62^. Proteins identified with the same set of peptides were automatically grouped together.

#### Protein quantitation and annotation

Two groups were compared, salmonids (Rainbow trout, Atlantic salmon, Brown trout) and non-salmonid teleosts (Goldfish, Common carp, Pikeperch). Only proteins identified in all the replicates of a same species with a minimum of 2 peptides were kept for further analysis. Genes from each species with a common ancestor (post teleost-specific whole genome duplication, TGD) were identified using Genomicus (GenomicusFish v04.02 - Gene Search) ^30^. Common ancestors were then used to perform inter-species protein comparison. For each protein, the mean number of spectral counts was calculated and normalized to the size of the protein (N-SC). If several proteins had the same common ancestor, the sum of their N-SCs was used for the quantification. The proteomic quantitative and statistical analysis was performed using ProStaR v1.28.0 ^63^. First, proteins not identified in at least 2 species of one of the two groups were filtered out. Then, data were normalized by centering N-SC distributions on the median. Missing values were inferred as follows: partially observed values (POV) were inferred using SLSA algorithm and values missing on entire condition (MEC) were inferred using DetQuantile method. Between-groups comparisons were performed with a limma test and Benjamini-Hochberg p-value correction. A –log10 (p-value) cut-off of 2.35 (p-value = 0.00447) was calculated for a FDR<1%.

Gene ontology analysis was performed using ShinyGO ^64^. First, human orthologs of the proteins identified in the rainbow trout coelomic fluid were obtained using genomicus. (<u>GenomicusFish v04.02 - Gene Search</u>). The corresponding human Ensembl gene numbers were then used for gene ontology annotation. The 500 most significantly enriched biological processes GO-terms (FDR < 0.05) were retrieved and hierarchically clustered. GO-terms sharing 70 % of their proteins were clustered together. Proteins of the main clusters, composed of at least 4 GO-terms, were further subjected to GO annotation to identify the high-level GO categories representing the different clusters, and the most represented GO-term in each cluster.

### Statistics

Statistical analyzes were performed using R software. Normal distribution of the variables was evaluated using Shapiro-Wilk test. When the distribution was normal, statistical difference of the means was evaluated using a t-test. Otherwise, a Wilcoxon - Mann Whitney test was performed.

## Ethical statement

Experiments and procedures were fully compliant with French and European animal welfare policies and followed guidelines of the INRAE LPGP Institutional Animal Care and Use Ethical Committee, which specifically approved this study.

## Data availability

All mass spectrometry proteomics data have been deposited on the ProteomeXchange Consortium (http://proteomecentral.proteomexchange.org) via the PRIDE partner repository ^65^ under the dataset identifier PXD034989 and 10.6019/PXD034989.

## Acknowledgements

The authors thank the INRAE ISC-LPGP for fish breeding. This study was funded by Agence Nationale de Recherche (ANR) grant EggPreserve # ANR-16-CE20-0001) to JB and CP.

